# The modified elevated gap interaction test: A novel paradigm to assess social preference

**DOI:** 10.1101/2023.10.30.564718

**Authors:** Chris I. De Zeeuw, Si-yang Yu, Jiawei Chen, Willem S. van Hoogstraten, Arn M.J.M. van den Maagdenberg, Laurens W.J. Bosman, Lieke Kros

## Abstract

Social deficits play a role in numerous psychiatric, neurological and neurodevelopmental disorders. Relating complex behavior, such as social interaction, to brain activity remains one of the biggest goals and challenges in neuroscience. Availability of standardized behavioral tests that assess social preference is however, limited. Here, we present a novel behavioral set-up and paradigm that we developed to measure social behavior, the modified elevated gap interaction test (MEGIT). In this test, animals are placed on one of two elevated platforms separated by a gap, in which they can engage in whisker-interaction with either a conspecific or an object. This allows quantification of social preference in real interaction rather than just proximity and forms an ideal setup for social behavior-related neuronal recordings. We provide a detailed description of the paradigm and its highly reliable, deep-learning based analysis, and show test results obtained from wild-type animals as well as genetic mouse models for disorders characterized by either hyposocial (autism spectrum disorder; ASD) or hypersocial (Williams Beuren syndrome; WBS) behavior. Wild-type animals show a clear preference for whisker interaction with another mouse rather than an inanimate object. This preference proved to be significantly smaller in mice carrying a mutation that can cause ASD in humans, whereas it is larger in WBS murine models. Differences in social preference became even more pronounced when corrected for hyperactive or repetitive behavior. The results indicate that MEGIT is a sensitive and reliable test for detecting and investigating social phenotypes.

## Introduction

Social behavior is crucial for survival and wellbeing, and social deficits play a role in numerous psychiatric, neurological and neurodevelopmental disorders^1-3^. Relating complex behavior, such as social interaction, to brain activity remains one of the biggest goals and challenges in neuroscience^2,4^. This topic is even more relevant if one considers how poorly we understand how changes at the molecular, cellular and network level lead to disorders involving social behavior such as autism spectrum disorder (ASD). Given the ethical and technical challenges in studying the mechanisms underlying ASD and other behavioral disorders, mouse models play an important role in elucidating the neuronal pathways and networks underlying social interaction in health and disease^5,6^. To this end, many mouse models harboring genetic mutations that are correlated to specific disorders in humans have been generated^5^. To validate these mouse models, animals are subjected to behavioral assays to establish phenotypes that resemble the human condition^5,6^. To characterize social behavior in mice, a variety of standardized, reproducible tests exist for the quantification of dominance, vocalizations or social preference^6-8^. Most translational studies, however, rely mainly on one of these tests, the three chamber test (TCT), which investigates social proximity rather than quality of social interaction^9,10^.

Mice are social animals engaging in a wide variety of social behaviors, including sniffing and whisker interaction^6,11,12^. These behaviors are, however, not quantified in the TCT. The standard TCT consists of a rectangular box divided into three chambers separated by small openings. One of the chambers holds a mouse under a wire-mesh cup, whereas the other holds an object. The test mouse can walk around freely, and preference for spending time in the social chamber is quantified^9-11^. The test is well-suited to analyze the quantity of social proximity, but does not assess actual social interaction. Alternatively, the gap paradigm can be used, in which mice are placed on elevated platforms separated by a gap in which social interactions can be monitored with video cameras^13^. Unlike the TCT, the classical gap paradigm has not been used as a standardized test to quantify aberrant social behavior. We, therefore, aimed to develop an alternative gap test that is as easy to perform as the TCT, but that provides more insight in the quality of the social interactions, while allowing quantification of specific related social parameters in an automated fashion. As such, this test, which we refer to as the modified elevated gap interaction test (MEGIT), can serve as a standardized paradigm to assess social preference in terms of actual interaction. Like the classical gap paradigm, the MEGIT consists of two elevated platforms; on one of them a test mouse is placed, while on the other platform, which is separated by a gap from the one with the test mouse, either another mouse or an inanimate object is placed. This allows investigation of social preference and the quality of social interaction in real encounters rather than just proximity. In addition, by tracking the test mouse, the relation between social phenotypes and often occurring comorbidities, like hyperactivity or repetitive behavior, can be studied. Finally, although not part of the current study, MEGIT also allows for an easy-to-use combination with electrophysiological recordings, as the behavioral setup does not require any physical structures above the head that may hinder connecting wires.

We first provide a detailed description of the MEGIT, its outcome measures, and our highly-reliable method of deep-learning based analysis using a specialized OptiFlex model^14^. We then show MEGIT results obtained from neurotypical wild-type animals, which form a basis for “normal” social behavioral patterns and preferences seen in this paradigm. To assess specificity and sensitivity of the MEGIT to detect social phenotypes, we furthermore set out to test mouse models for disorders characterized by either hyposocial (ASD) or hypersocial (Williams Beuren syndrome; WBS) behavior.

ASD is a highly prevalent, neurodevelopmental disorder characterized by social deficits, ranging from a complete lack of interaction to difficulty understanding sarcasm and repetitive behavior^3^. Various mutations have been associated with the disorder, including several mutations in postsynaptic density proteins, such as *Shank2*^15-17^. Global deletion of *Shank2* has been shown to result in a pronounced ASD-like phenotype in mice including decreased social preference and hyperactive/repetitive behavior^18-20^. Conversely, WBS is a rare syndrome caused by a hemizygous deletion of 25 to 27 genes on chromosome 7q11.23, resulting in both cardiovascular and neuropsychological symptoms^21-24^. Patients with WBS often have reduced stranger awareness, leading to a strong tendency to engage with other people, a trait often summarized as hypersociability^23-27^. Mouse models for WSB, harboring deletion of genes on chromosome 5, corresponding to human chromosome 7, mimic the symptoms seen in humans including hypersociability^28-31^. Because of their opposing social phenotypes, these two mouse models were selected to assess sensitivity of the MEGIT to detect differences in social preference.

## Materials and Methods

### Animals

Data were obtained from adult (8-20 weeks) mice that were confronted with juvenile (22-31 days) mice of either sex. All mice had a C57BL/6J background (Charles River), while in specific experiments germline *Shank2*^-/-18,20^, or WBS^32^ mice were used. Mutant lines were bred in-house and compared to their wild-type littermates. To generate the WBS mouse model, CRISPR/Cas9 technology was used to remove the genes between *Ncf1* and *Fkbp6* on mouse chromosome 5, corresponding to the same genes on chromosome 7q11.23 in humans^32,33^. All mice were group housed in enriched cages, had ad libitum access to food and water and were held on a 12:12h light/dark cycle. Experiments were done in accordance with European, national and institutional laws and guidelines. Protocols were reviewed and approved by the institutional experimental animal committee.

### The MEGIT setup

The MEGIT setup is essentially a box made of clear Plexiglas with two platforms at a height of approximately 20 cm, lit from underneath by an LED-screen (Fig 1A). The distance between the platforms is adjustable ensure that mice of any size are able to interact in the gap between the platforms, but cannot jump from one to the other. Two adjustable cameras (Basler AG, Ahrensburg, Germany) are mounted above the setup to record the behavior of the animals. One records at 60 Hz and captures both platforms and overall behavior (overview video) whereas the other zooms in on the gap to allow detailed assessment of interactions at a higher rate (100 Hz; zoom video; Fig 1A, B). Recordings were made and synchronized using LabVIEW software (National Instruments, Austin, TX).

**Fig 1.**
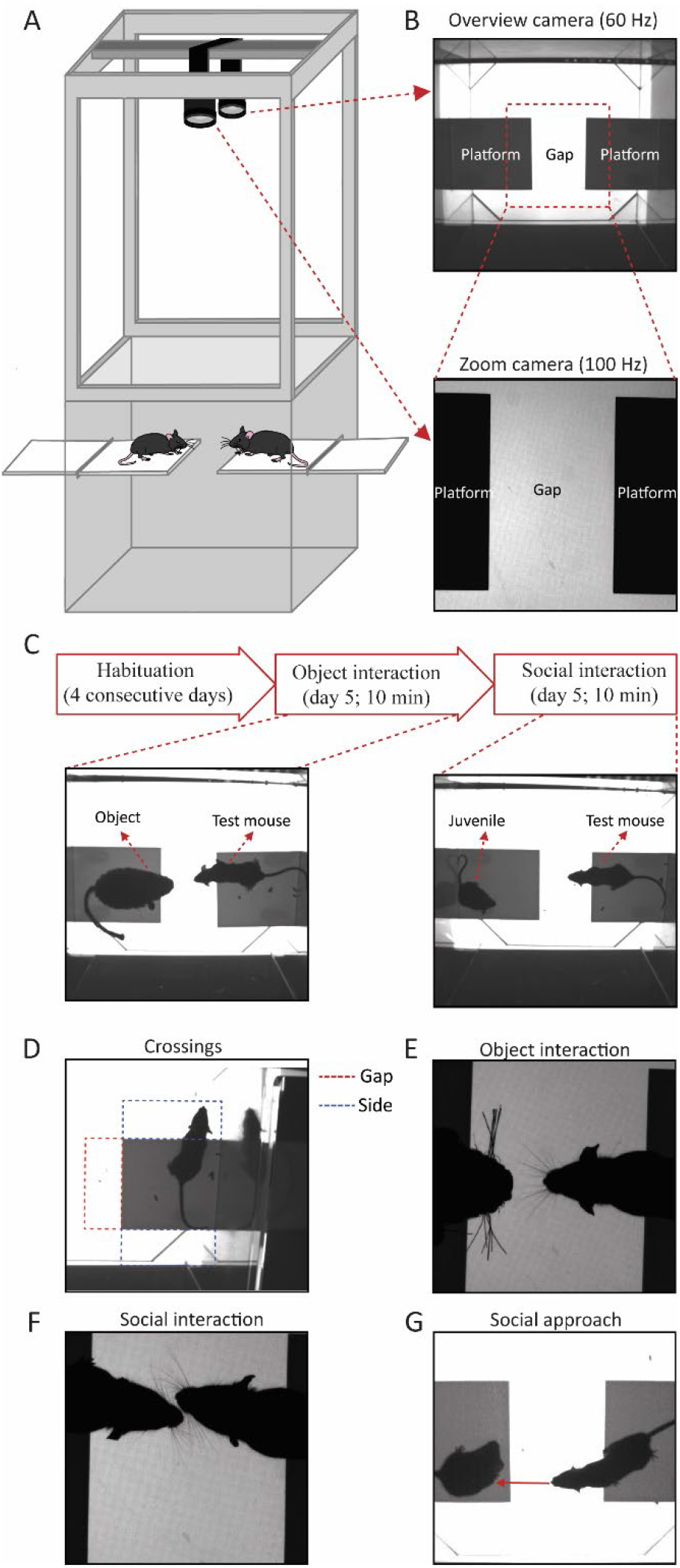
The MEGIT explained. **A**. Schematic drawing of the MEGIT setup consisting of a box with two adjustable platforms on which the mice can be placed and 2 cameras capturing the behavior. **B**. Examples of images captured by an overview camera (top; 60Hz) used to assess overall behavior and a zoom camera (bottom; 100 Hz) specifically aimed at the gap; the area in between the platforms where interaction takes place. **C**. Experimental design. All mice are habituated to the setup for 4 consecutive days before the test, consisting of 10 min of object interaction (left bottom picture) and 10 min of social interaction (right bottom picture), takes place on day 5. **D-G**. Representative examples of the outcome measures; crossings (**D**), object interaction (**E**), social interaction (**F**) and social approach (**G**). Number of occurrences, time spent and average duration is calculated for each of these measures.

### Experimental design of the MEGIT

Experimenters were blinded to genotype until completion of all analyses. Animals were first habituated to both the experimenter and the setup to reduce stress. Habituation was performed over four consecutive days for increasing amounts of time (5-20 min) without another mouse or object present on the opposing platform, after which the experiment took place on day five (Fig 1C). After starting recording, the test mouse was placed on one of the platforms and the object on the other. Behavior was recorded for 10 minutes. In case the mouse jumped off the platform, the recorded time was increased by ~30 s to allow for the removal of frames without the animal being on the platform. This ensured that the total length of the videos after preprocessing was no less than 10 min. After completion of the object video, a juvenile wild-type mouse was placed on the other platform and a total of 10 min of social behavior was recorded. Juvenile mice were used for the social videos to avoid either aggressive or sexual behavior from a male test mouse. Upon completion of the two videos, the setup was thoroughly cleaned with 70% ethanol.

### MEGIT outcome parameters

Several behavioral parameters were used to compare both the degree of repetitive/hyperactive behavior and social preference between mice and genotypes. To assess repetitive/hyperactive behavior, crossings were quantified. A crossing is an instance where an animal sticks its head outside the platform in a predefined region of interest (ROI). For it to be included as a crossing, both ears had to fully surpass the platform for at least six consecutive frames (~100 ms) in the overview video. Four ROIs were defined, the gap (Fig 1D; red dashed line) and the sides (Fig 1D; blue dashed lines) for test mice and just the gap for juvenile mice. If the test animal’s head was in one of the 4 corners outside the platform; e.g., between the gap and one of the sides or one of the sides and the back wall, it was only included as a crossing if >50% of the head was in an ROI and both ears surpassed the extrapolated border between ROI and platform.

Gap crossings were subsequently used to select and quantify object interactions and social behavior. For object and social interactions, frames of the zoom videos corresponding to gap crossing containing frames in the overview video were inspected. An interaction was defined as an instance where the whiskers of the test mouse touches either the object or the juvenile mouse (Fig 1E, F). Social interaction can occur when only one of the two animals has its head completely in the gap. Because the animal can interact with the object whenever it prefers but can only socially interact when both animals choose to do so, social approach was included as an additional measure for social behavior. This accounts for the option that the test mouse approaches the juvenile mouse, but the juvenile mouse does not react. Social approaches were selected from the overview video and were defined as instances where the test mouse reaches for the juvenile mouse during a gap crossing for more than six consecutive frames. An event was included as reaching when a virtual line, drawn from the middle of the head through the nose, touches the juvenile animal (Fig 1G). For all these parameters, the number of occurrences/min, time spent and the average duration was calculated.

To correct for the effects of potential hyperactive / repetitive behavior on interactions and most notably approaches, raw interaction and approach data was normalized to gap crossings. This was done by dividing each interaction / approach variable by the corresponding gap crossing variable of the same video. For example, normalization of the number social approaches to gap crossings was done by dividing the number of approaches by the number of gap crossings by the test mouse in the social video. This allows for a measure of proportion of gap crossings containing a social approach.

### MEGIT analyses

To efficiently and objectively analyze the videos, a specialized OptiFlex model^14^ was used to automatically label and track both ears and the nose of each animal. The model was trained by feeding it 5326 frames with the ears and nose manually labeled by two independent observers (Fig 2A) after which it was able to provide the trajectory of the animal’s head (Fig 2B). A custom-written program (Python; Python Software Foundation, Wilmington, DE) was used to streamline the analysis process. First, each frame of both videos was labeled with consecutive numbers to allow for optimal synchronization of the two videos after removing excess frames. Excess frames consisted either of frames where the mouse was not on the platform (at experiment initiation or upon a jump off the platform) or frames that exceeded the total time of 10 minutes and were removed after labeling. Next, the ROIs were defined manually. OptiFlex tracked trajectories of the nose and both ears were subsequently used to identify crossings. For this, all three markers had to have crossed the platform-side border of an ROI and the area of the triangle formed by those three markers had to be in an ROI > 50%. Only frames meeting these criteria were classified as instances where the animal crossed into a specific ROI.

**Fig 2.**
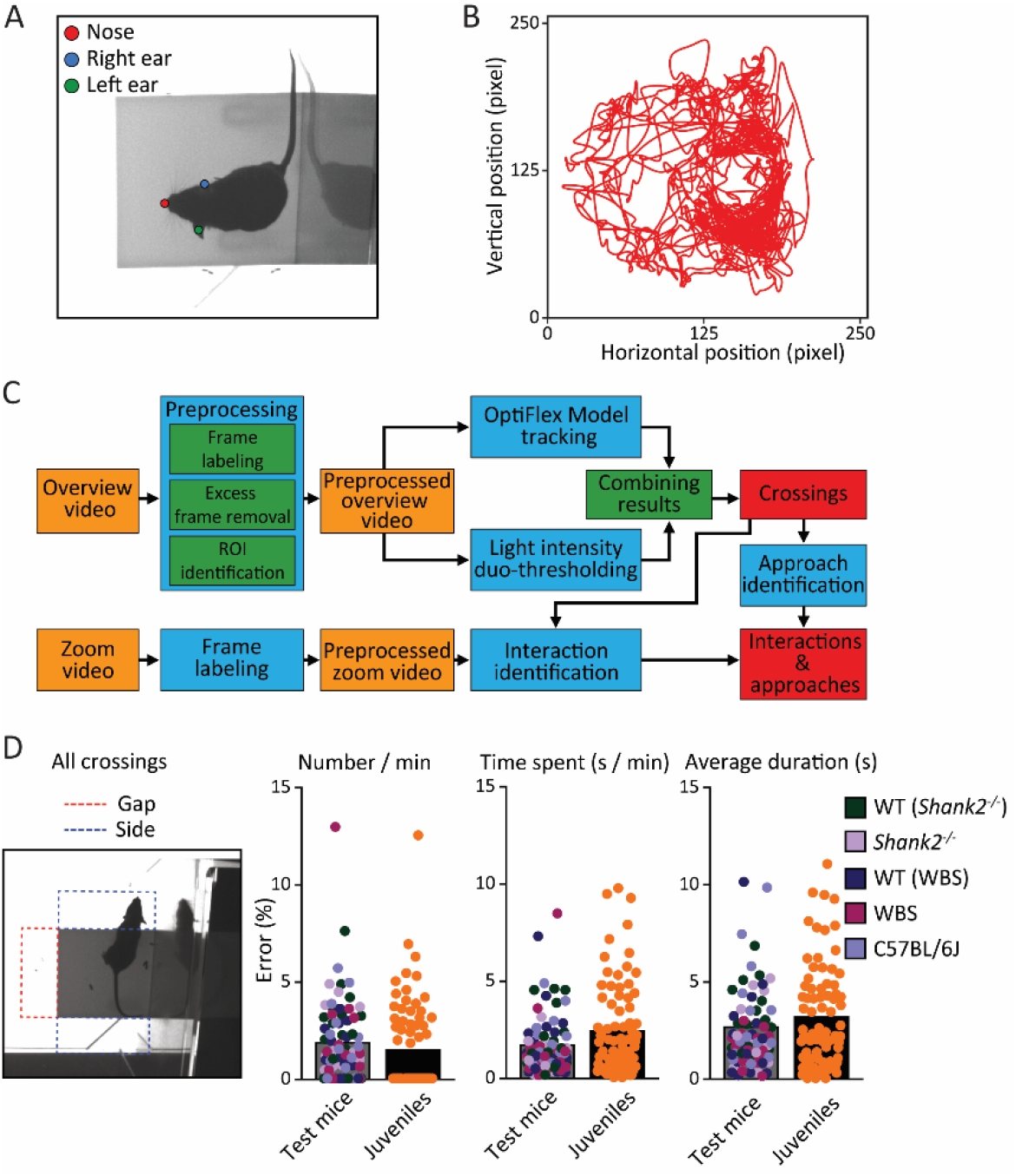
Machine learning based MEGIT analysis. **A**. Example of a frame in which the nose and ears of the test mouse are labeled. Labeled frames were used to train the OptiFlex model to track the position of the head of both the test mouse and juvenile throughout the video’s. **B**. Representative example of the trajectory of the nose over one 10 min video. **C**. Schematic representation of the analysis process. First, preprocessing is done semi-manually, followed by OptiFlex model based tracking of the position of the head to allow for the determination of the start and end of crossings. Subsequently, interactions and approaches are manually annotated based on selection of frames containing gap crossings. **D**. Quantification of the accuracy of the automated detection of crossings. All crossings (left picture; both video’s; gap crossings (red) and side crossings (blue)) of all 76 test mice and juveniles used for this study were annotated both manually and using the OptiFlex model. Bar plots show the error rate for each individual mouse for the total number of crossings per minute (left), total time spent (middle) and average duration (right). Grey bars show data of test mice (genotypes color coded) and black bars with orange dots show the corresponding juvenile data. Average error rates are well below 95%.

In addition to the model classification, a traditional computer vision method was employed to further refine the classification results. For each ROI, the mean light intensity throughout the video was computed. Subsequently, two distinct light intensity thresholds were defined for each region based on the size of the animal: “the greedy threshold”, which focused on minimizing false-negative crossings, and “the conservative threshold”, which aimed at minimizing false-positive crossings. Binary results were then generated using both thresholds. To enhance accuracy, the classification results obtained from the model predictions were subjected to a stringent filtering process: negatives from the greedy detection were forcefully rejected, while positives from the conservative detection were emphatically accepted.

Finally, a straightforward sliding window algorithm was applied to detect crossings. The start of a crossing was defined as the first frame of at least six consecutive frames showing a crossing in an ROI, whereas the end was set as the first of at least six consecutive frames without a crossing in the ROI. Once the crossings were detected, the algorithm proceeds to calculate the number of crossings, the time spent and the average duration for each ROI. The automatically detected gap crossings were subsequently used to select frames in which potential interactions and approaches could be manually identified (Fig 2C). For this study, crossings from all test mice (N=76) and juveniles (N=15) were both manually and automatically analyzed to assess the reliability of the automatized process. An average accuracy of >95% was reached for all variables (Fig 2D).

### The Three Chamber Test (TCT)

Mice that were tested using the MEGIT were also subjected to the TCT. The setup consists of a rectangular 130 × 80 × 80 cm, clear Plexiglas box, divided into three chambers separated by two doors. Behavior was continuously recorded using a 25 Hz camera mounted above the setup using the open-source software Bonsai (https://bonsai-rx.org). Before test initiation, the test mouse was habituated to the middle chamber for 5 minutes and subsequently allowed to explore all chambers of an empty setup for 10 minutes. After this, the test-mouse was confined to the middle chamber and a habituated sex- and age-matched wild-type mouse was placed under a cup in one random outer chamber and an empty cup in the other. The doors separating the chambers were then opened and the behavior of the test mouse was recorded for 10 min. Number of entries and time spent in the individual chambers were subsequently calculated using the open-source software OptiMouse^34^.

### Statistical analyses

Differences between outcome parameters within the pooled wild-type data were tested using paired-samples *t* tests and Bonferroni corrected *p-*values. Differences between genotypes were assessed using non-parametric Mann-Whitney U tests because assumptions of normality of distributions and equality of variances were often violated. Correlations between MEGIT outcome parameters and TCT results were tested using Pearson correlations. All statistical analyses were performed using SPSS 27 software (IBM, Armonk, NY) using two-tailed testing.

## Results

### Social preference in wild-type animals

In order to assess normal, non-mutant behavior in the MEGIT, we first analyzed the pooled data of all wild-type groups (*Shank2*^-/-^ wild-type littermates, WBS wild-type littermates and C57BL/6J mice without mutant littermates). To test whether a test mouse preferred gap crossings when another mouse was present on the other platform over other, non-social crossings, we first quantified the different types of crossings (Fig 3A). The number of social gap crossings was significantly higher than both the number of object gap crossings and the number of side crossings (*p*<0.001 for both; Fig 3B, C). In addition, whereas the number of social gap crossings was significantly higher than the number of side crossings in the same epoch (*p*<0.001), such difference could not be found in the object video (Fig 3C). Similarly, both time spent on and average duration of social gap crossings were significantly higher for social gap crossings as compared to gap crossings in the object video (*p*<0.001 for both; Fig 3B, C). These results indicate a clear preference for exploration of the gap when being across a conspecific, rather than an object.

**Fig 3.**
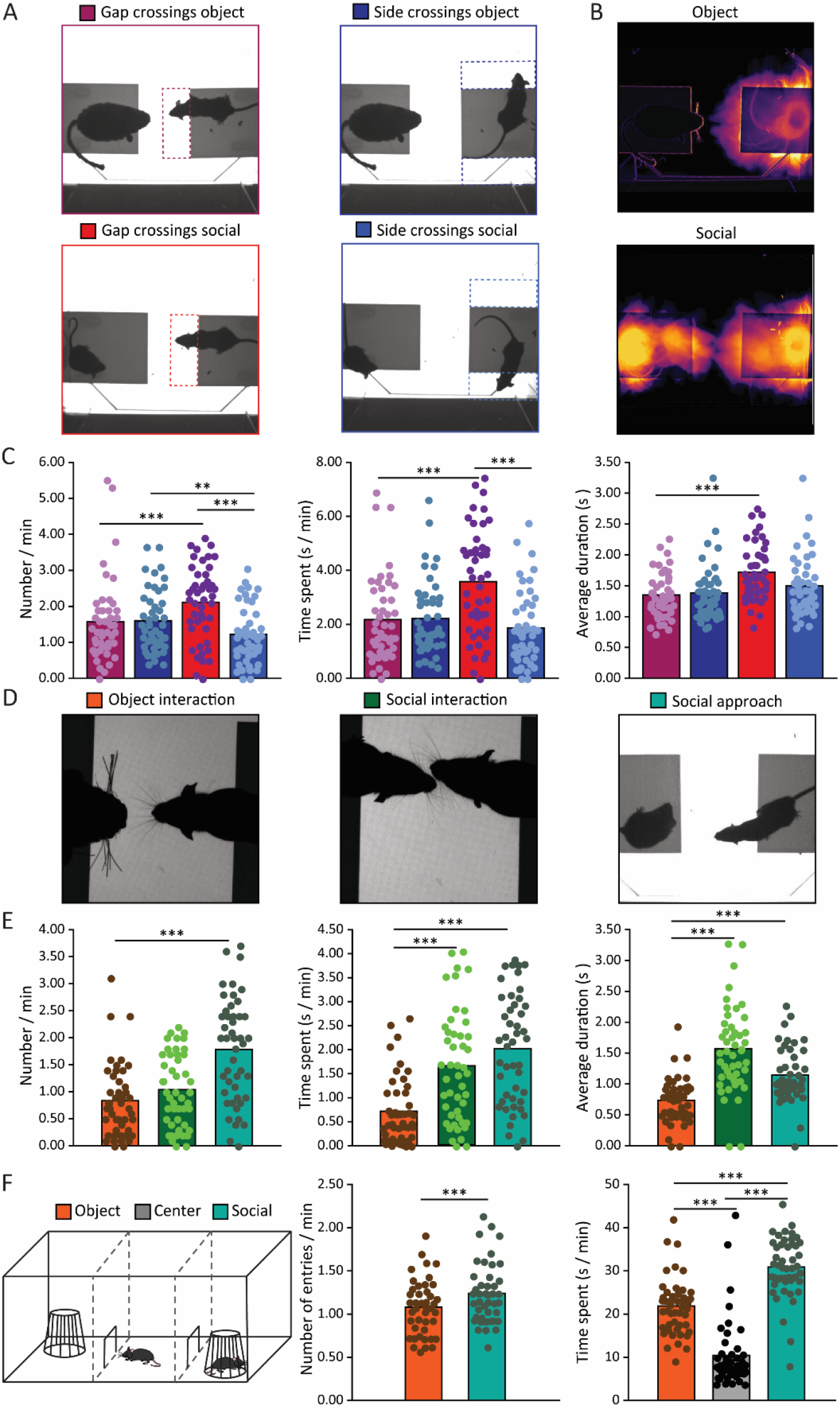
MEGIT results of wild-type animals. **A**. Representative examples of crossings during the object video (top) and social video (bottom) in the gap (left) or sides (right). **B**. Representative example of the standard deviation of light intensity of the object (top) and social video of one wild-type mouse. **C**. Quantification of the number of (left), time spent on (middle) and average duration (right) of gap (purple) and side (dark blue) crossings in the object video and gap (red) and side (light blue) crossings in the social video for all wild-type mice (N=47). **D**. Representative examples of object interaction (left), social interaction (middle) and social approach (right). **E**. Quantification of the number of (left), time spent on (middle) and average duration (right) of object interactions (orange), social interactions (dark green) and social approaches (light green) of all wild-type mice (N=47). Note that comparisons between social approaches and interactions were not made because of substantial overlap. **F**. Schematic representation of the three chamber test (left) and quantification of the number of entries (middle) and time spent (right) in the object chamber (orange), social chamber (green) or middle chamber (grey; only shown for time spent) of all wild-type mice (N=47). For all comparisons: ^*^*p* < 0.05, ^**^*p*<0.01, ^***^*p* < 0.001 (Paired-samples *t* tests).

Since not all gap crossings are directed towards the other mouse or the object, we subsequently studied social approaches and physical interactions with either the other mouse or the object (Fig 3D). Of these, the social approaches were the most frequent (*p*<0.001), while physical interactions of the other mouse and the object occurred roughly equally often (Fig 3E). However, social events, whether approaches or interactions, lasted markedly longer than object interactions (*p*<0.001 for both time spent and average duration; Fig 3E). Because data of the different wild-type groups was pooled, we compared the social data of the separate groups as a control and found no significant differences on any of the variables (Fig S1A, B).

Finally, all animals were subjected to the traditional TCT as well resulting in a significantly increased number of entries and time spent in the social chamber as compared to the object chamber (*p*<0.001 for both; Fig 3E). Together, these results show that wild-type animals exhibit a clear social preference on both the MEGIT and the canonically used TCT.

### Repetitive/hyperactive behavior in mouse models for ASD and WBS

A potential confounder for the interpretation of social behavior is the presence of repetitive or hyperactive behavior that could increase the overall number of crossings. Before evaluating social phenotypes of mutant mice, we, therefore, first focused on repetitive and hyperactive behavior in a mouse model for ASD with a known hyperactive phenotype, global *Shank2* knockout mice^18^, and a novel mouse model for WBS^32^, in comparison to their respective wild-type littermates. When comparing data on all crossings (gap and sides, both videos), *Shank2*^-/-^ mice showed a remarkable three-fold increase in number of crossings (*p*<0.001), a marked decrease in duration of crossings (*p*<0.001), and a milder increase in time spent on crossings (*p*<0.05; Fig 4A). When specifically comparing data on gap crossings during either the object video (Fig 4B) or the social video (Fig 4C), a similar pattern of increased number (*p*<0.001 for both), increased amount of time spent (*p*<0.001 and *p*<0.01 for object and social gap crossings respectively) and shorter average duration (*p*<0.01 and *p*<0.001 for object and social gap crossings respectively) of gap crossings was shown as compared to their wild-type littermates. Investigation of the same parameters but regarding side crossings during either the object video (Fig S2A) or the social video (Fig S2B) indicated a similar increase in number (*p*<0.001 for both) and decrease in average duration (*p*<0.001 for both) compared to control animals, but no differences in time spent. These data indicate that *Shank2*^-/-^ mice indeed show a pronounced hyperactive phenotype in the MEGIT.

**Fig 4.**
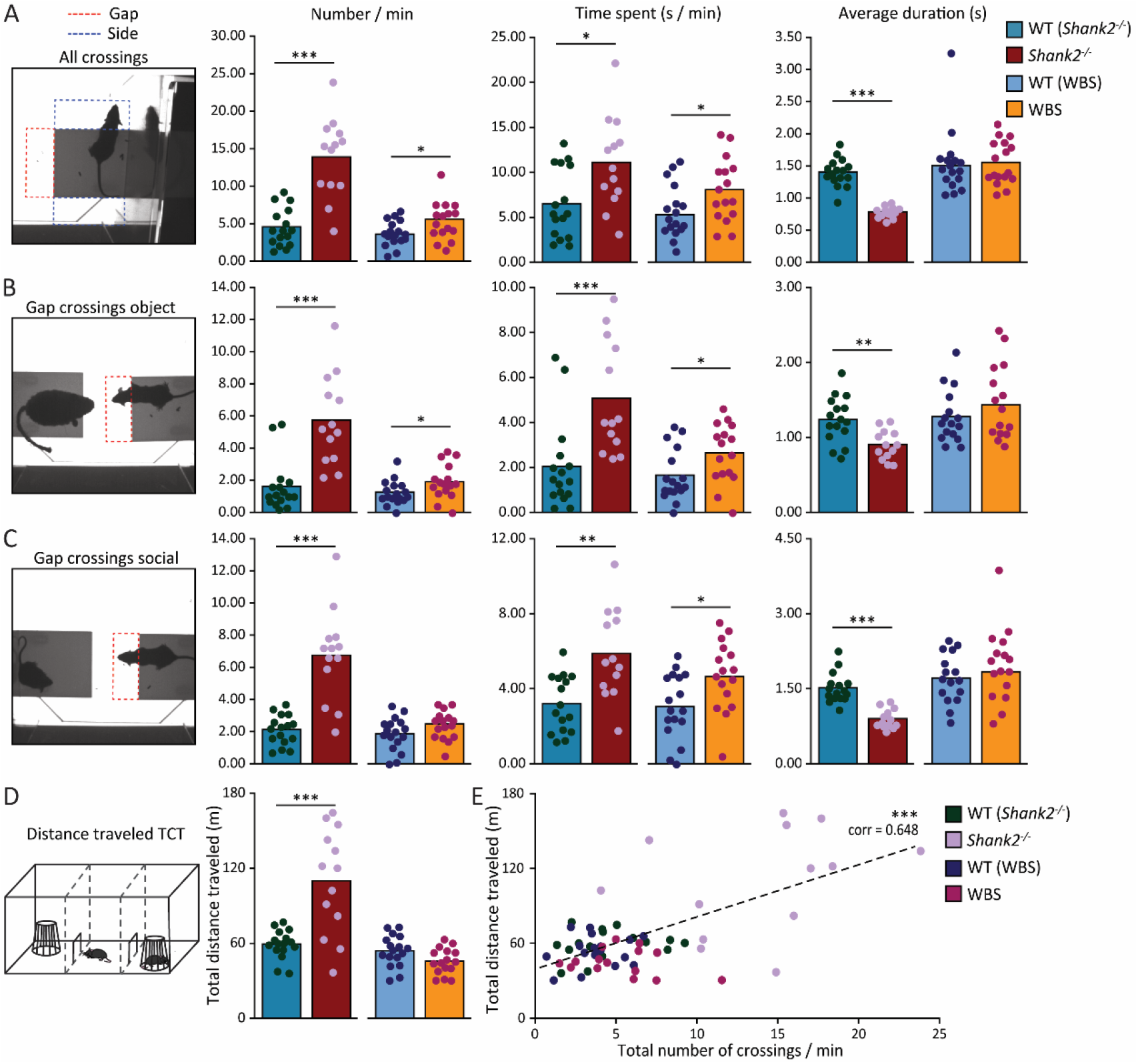
Assessment of hyperactive / repetitive phenotypes. **A**. Quantification of the number of (left), time spent (middle) and average duration of all crossings (all ROIs, both videos) for *Shank2*^-/-^ mice (dark red; N=13), their wild-type litter mates (green blue; N=16), WBS mice (orange; N=16) and their wild-type littermates (blue; N=17). **B**. As in **A**. but specifically for gap crossings in the object video. **C**. As in **A**. but specifically for gap crossings in the social video. **D**. Quantification of the distance traveled in the TCT. Color coding as in **A-C. E**. Correlation between distance traveled in the TCT and the total number of crossings in the MEGIT. Color coding as in **A-C**. For all comparisons: ^*^*p* < 0.05, ^**^*p*<0.01, ^***^*p* < 0.001 (Mann-Whitney U tests and Pearson correlation).

The same tests and analyses were performed in our cohort of WBS mice. WBS mice also show a hyperactive phenotype, albeit much less pronounced than *Shank2*^-/-^ mice. Comparisons of all crossings indicate a higher number of (*p<*0.05) and more time spent on (*p<*0.05) on crossing without a difference in duration (Fig 4A). Calculations on gap crossings demonstrate an increase in time spent for both object and social gap crossings (*p<*0.05 for both) and a higher number only for gap crossings in the object video (*p<*0.05; Fig 4B, C). No differences were observed for any of the parameters regarding side crossings.

Subsequently, the distance traveled was used as an estimate of hyperactive behavior in the TCT. Again, *Shank2*^-/-^ mice were much more active than their wild-type littermates (*p*<0.001), whereas WBS mutants were not significantly different from control mice (Fig 4D). Relating the total number of MEGIT crossings with distance traveled in the TCT produced a significant correlation (*p*<0.001) demonstrating the robustness of activity level in these animals.

### Social phenotypes in ASD and WBS mouse models

Next, the MEGIT was tested for its ability to identify changes in social behavior in the two different mouse models: *Shank2*^-/-^ mice that model hyposocial behavior^18-20^ and WBS mice that model hypersociability^24,27^. To this end, the differences between social variables and object interactions were quantified as a measure of social preference. *Shank2*^-/-^ mice showed a lower and WBS mice a larger social preference, in terms of number of interactions, than their respective wild-type littermates (*p<*0.001 and *p<*0.05 for *Shank2*^-/-^ and WBS mice, respectively; Fig 5A, Fig S3A, B). In WBS mice, the increased social preference was also reflected in the difference between social approaches and object interactions both in number of occurrences (*p<*0.001) and time spent (*p<*0.05), but such difference was absent in *Shank2*^-/-^ mice (Fig 5B, Fig S3A, C).

**Fig 5.**
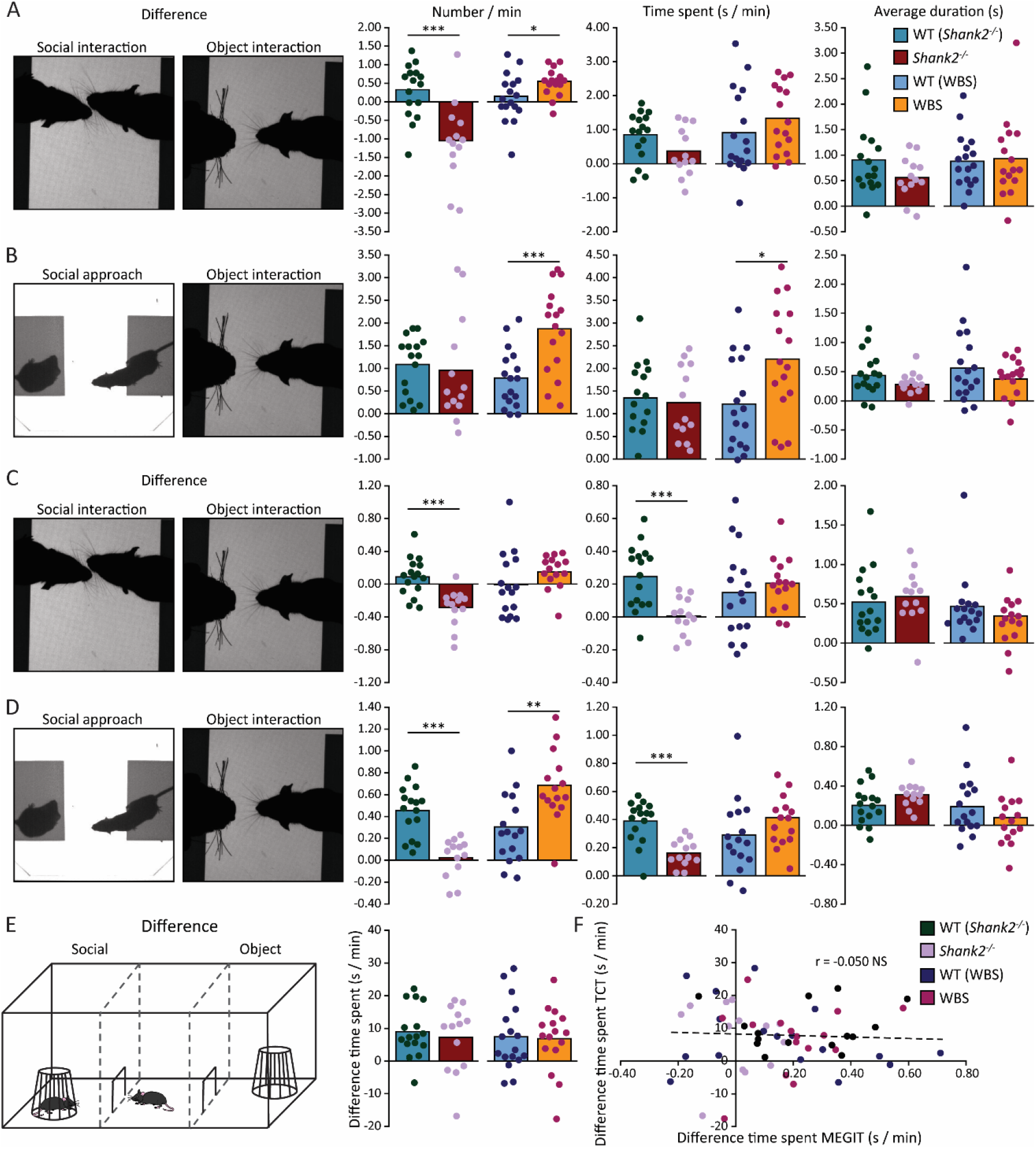
Assessment of phenotypes in social preference. **A**. Quantification of the difference between social and object interactions in number (left), time spent (middle) and average duration (right) for *Shank2*^-/-^ mice (dark red; N=13), their wild-type litter mates (green blue; N=16), WBS mice (orange; N=16) and their wild-type littermates (blue; N=17). **B**. As in **A**. but for the difference between social approaches and object interactions. **C, D**. As in **A, B**. but for differences between social interactions (**C**) or social approaches (**D**) and object interactions after normalization to gap crossings. **E**. Quantification of differences in time spent between the social chamber and object chamber (right) of the TCT (right). Color coding as in **A-D. F**. Correlation between differences in time spent between the social chamber and object chamber in the TCT and differences between social and object interactions in the MEGIT. Color coding as in **A-C**. For all comparisons: ^*^*p* < 0.05, ^**^*p*<0.01, ^***^*p* < 0.001 (Mann-Whitney U tests and Pearson correlation).

Interpretation of the results is obfuscated by differences in activity level. Particularly *Shank2*^-/-^ mice show a pronounced degree of hyperactivity and are roughly three times more likely to engage in gap crossings than their wild-type littermates. As a consequence, an increased number of social approaches can be expected in *Shank2*^-/-^ mice. To isolate the impact of the social context from hyperactivity, the interaction and approach data were normalized to gap crossings, thus creating an estimate for the proportion of gap crossings dedicated to either social behavior or object interactions. When normalized to gap crossing data, *Shank2*^-/-^ mice show a significantly lower preference for social interaction than their wild-type littermates, both in number of instances and in time spent (*p*<0.001 for both) whereas no differences between WBS mice and wild-type controls were observed (Fig 5C, Fig S3D, E).

After normalization to gap crossings, pronounced phenotypes emerged: where WBS mice show a significant social preference in number of social approaches versus object interactions (*p*<0.01), *Shank2*^-/-^ mice show a significantly lower social preference compared to wild-types, both for the number of occurrences and time spent (*p*<0.001 for both; Fig 5D, Fig S3D, F).

Results from the same animals subjected to the TCT failed to show any significant differences. Moreover, all difference scores between time spent in the social chamber and time spent in the object chamber were positive, indicating social preference (Fig 5E). Unsurprisingly, no significant correlations were found between social preference scores in the MEGIT and the TCT (Fig 5F). This indicates that the MEGIT is a sensitive paradigm to test for both hypo- and hypersocial phenotypes, especially when data are corrected for hyperactive behavior by normalizing to gap crossings.

## Discussion

In this study, we present the MEGIT, a novel standardized test to quantify social preference by assessing real social interaction. MEGIT consists of two elevated platforms separated by a gap. On one platform, the test mouse is placed and interactions of this mouse with either an object or a conspecific across the gap are recorded. Analysis of the videos can be done reliably, objectively and efficiently using a specialized OptiFlex model^14^. As mice are social animals, they have an innate interest in other mice^35-37^, and this is well-reflected by our observation that wild-type mice increase the time they are spending in the gap when another mouse is present on the opposite platform. Indeed, wild-type mice exhibit a clear preference for engaging with another mouse rather than with an inanimate object. In MEGIT, this is quantified as an increase in the durations of social interaction and social approach. To test whether MEGIT is sensitive enough to reliably detect phenotypes in social behavior, two genetic mouse models with known alterations in their social behavior^18,20,28,32^ were subsequently tested. In line with the behavior typically observed in humans carrying these mutations, we found hypersocial behavior in WBS mice, whereas *Shank2*^-/-^ mice showed a markedly decreased social preference. Automated analysis of MEGIT was shown to be able to discriminate between social behavior and potential confounders, such as hyperactivity. Altogether, these results indicate that MEGIT is a sensitive and reliable test that can be used to detect social phenotypes.

In MEGIT, wild-type mice engage equally often with an object than with another mouse, but the social interactions last longer than those with the object. This indicates that social interactions indeed follow a different strategy than non-social interactions. However, the interpretation of these findings is complicated, as there is a difference in the availability of the object and of the other mouse. The other mouse, obviously, can choose to make itself available – or not -, while the object is always there. Despite this dependency on the behavior of the other mouse, non-mutant test mice still spend more time on social interaction. To nevertheless control for the social behavior of the other mouse, we sought to include a measure for social preference that was solely dependent on the test mouse. To this end, social approaches were included as an additional outcome measure. Approaches are initiated by the test mouse, irrespective of whether the approaches are reciprocated by the second mouse. It is possible that the test mouse reaches towards the other mouse by chance rather than due to intentional social approach. However, *Shank2*^-/-^ and WBS mice show clear differences with their respective wild-type littermates indicating the sensitivity of MEGIT to detect phenotypes in either direction.

Social behavior is extremely complex and rodents can engage in a wide variety of social behaviors other than facial touch and social approach as is assessed with MEGIT^11^. Technical advances and sophisticated tracking methods are making it increasingly feasible to assess the full spectrum and complexity of social behaviors^38,39^. This is a valuable development that should be used to study complex social phenotypes in rodent models. Pursuing that avenue does however not negate the need for more standardized, easy-to-interpret social paradigms. Whereas standardized tests are less reflective of the full complexity of behaviors, they can be more useful and robust in testing specific traits characteristic of disorders such as social preference or repetitive behavior^35^. In addition, social interaction in a home cage setting can occur because a test mouse chooses to engage. However, avoiding cage mates is difficult and it is feasible that some interactions happen without the test animal wanting to engage, complicating data interpretation. MEGIT allows for real interaction only if mice choose to do so. Furthermore, electrophysiological recordings during social and object interaction can be performed in the MEGIT^13,40-43^. By doing so, neuronal responses can be triggered by a specific type of social behavior and compared to essentially the same behavior minus the social aspect. This can potentially lead to a better understanding of the contribution of social rather than the sensory and motor aspects to neuronal responses. We therefore advocate investigating social behavior and phenotypes by both studying complex home cage behavior and using standardized tests including both TCT and MEGIT.

## Acknowledgements

Financial support was provided by ZonMw Veni (91619109; L.K.), Erasmus Medical Center Fellowship (L.K.), the Medical Delta Programme (MD 01092019-31082023; L.K., C.I.D.Z. & A.M.J.M.V.D.M), LSH-NWO Crossover (INTENSE TTW/00798883; L.K. & C.I.D.Z.), by Health Holland to promote public private partnerships (TKI-LSH EMCLSH21017: L.W.J.B.), NWO-ALW (824.02.001; C.I.D.Z.), ZonMw-TOP (91120067; C.I.D.Z.), ERC-advanced (GA-294775 C.I.D.Z.) and NWO-Gravitation Program (DBI2; C.I.D.Z.). We thank Dr. Aleksandra Badura for kindly allowing us to use her TCT setup.

## Figure legends

**Fig S1.**
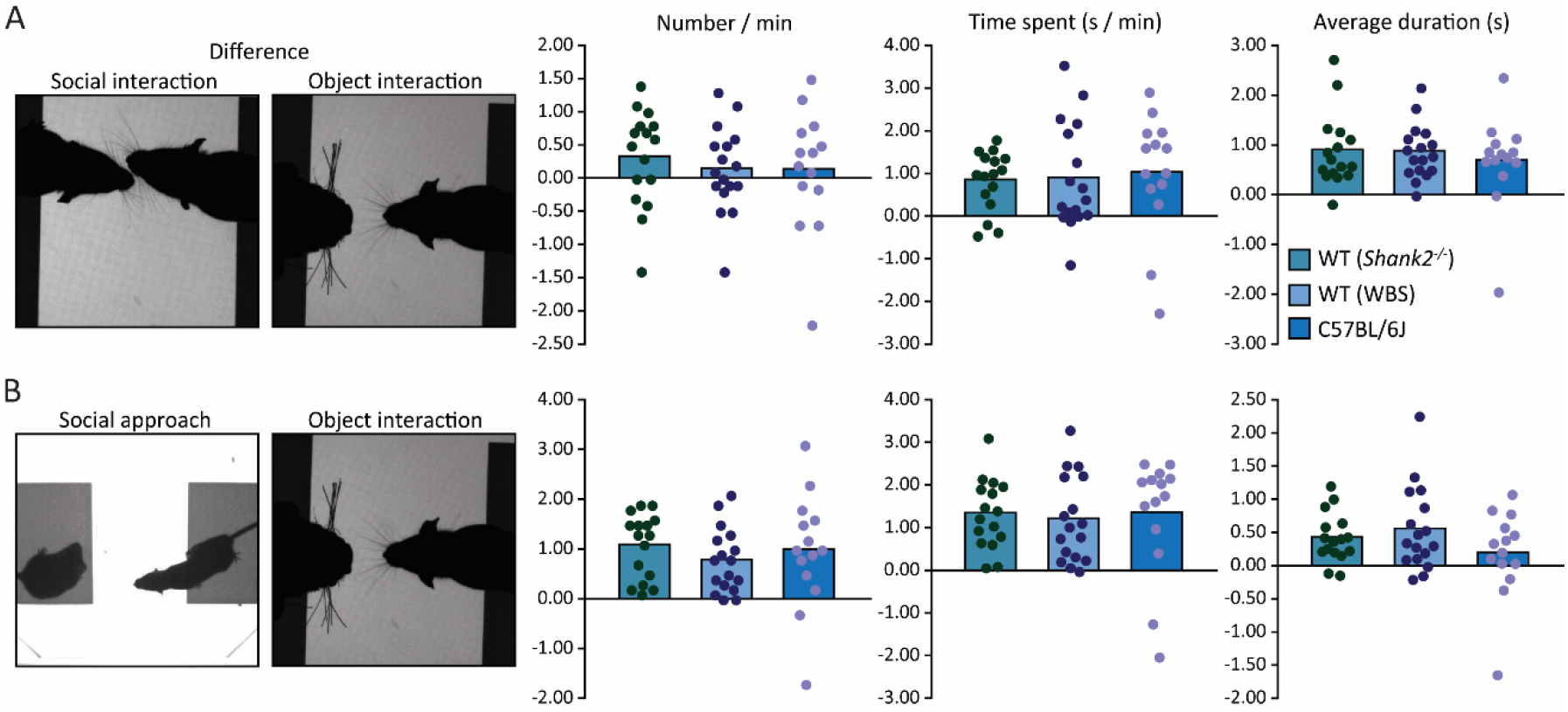
Comparisons of social preference in the different wild-type groups. **A**. Quantification of the difference between social and object interactions in number (left), time spent (middle) and average duration (right) for *Shank2*^-/-^ wild-type littermates (green blue; N=16), WBS wild-type littermates (blue; N=17) and C57BL/6J mice without mutant littermates (darker blue; N=14). All comparisons were non-significant (Mann-Whitney U tests).

**Fig S2.**
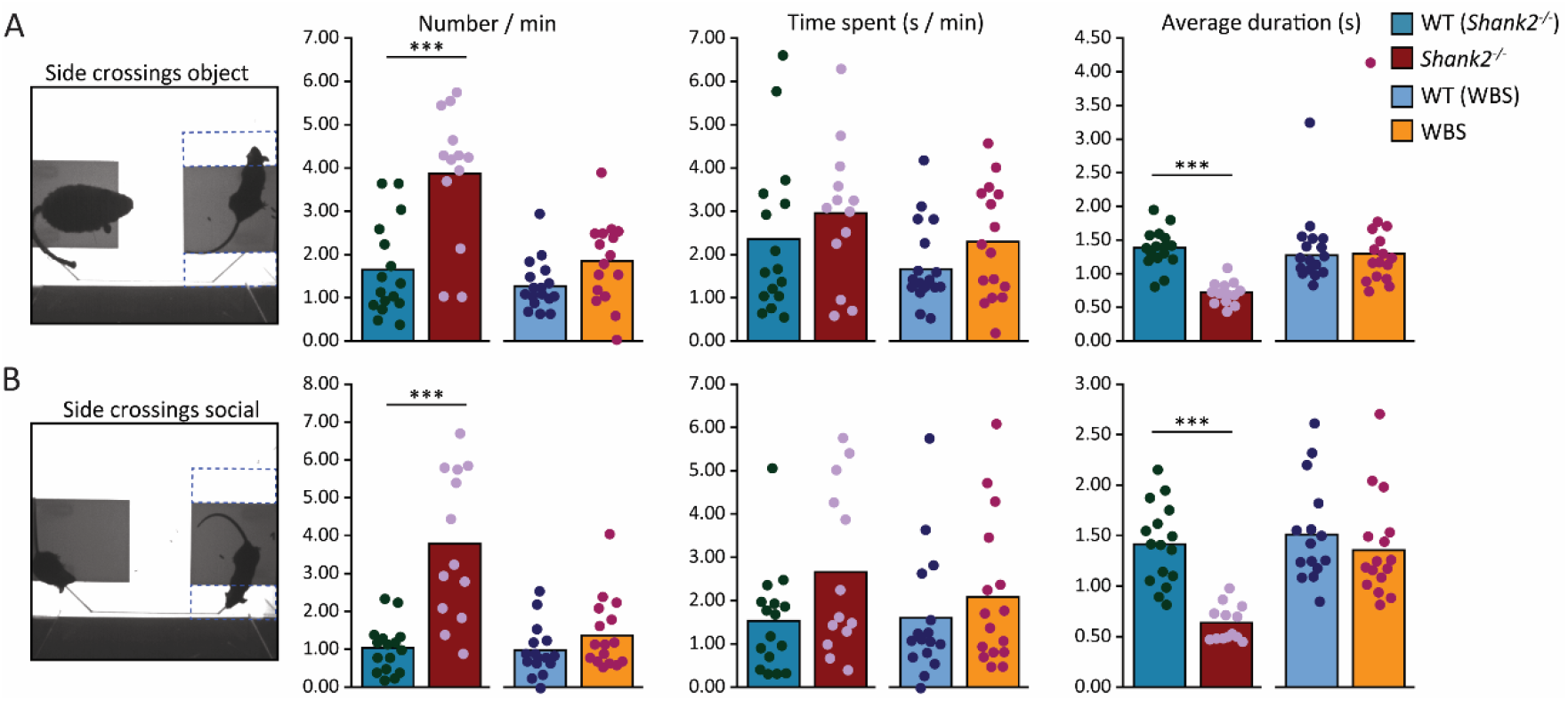
Assessment of hyperactive / repetitive phenotypes (side crossings; extension of Fig. 4). **A**. Quantification of the number of (left), time spent (middle) and average duration of side crossings in the object video for *Shank2*^-/-^ mice (dark red; N=13), their wild-type litter mates (green blue; N=16), WBS mice (orange; N=16) and their wild-type littermates (blue; N=17). **B**. As in **A**. but for side crossings in the social video. For all comparisons: ^*^*p* < 0.05, ^**^*p*<0.01, ^***^*p* < 0.001 (Mann-Whitney U tests).

**Fig S3.**
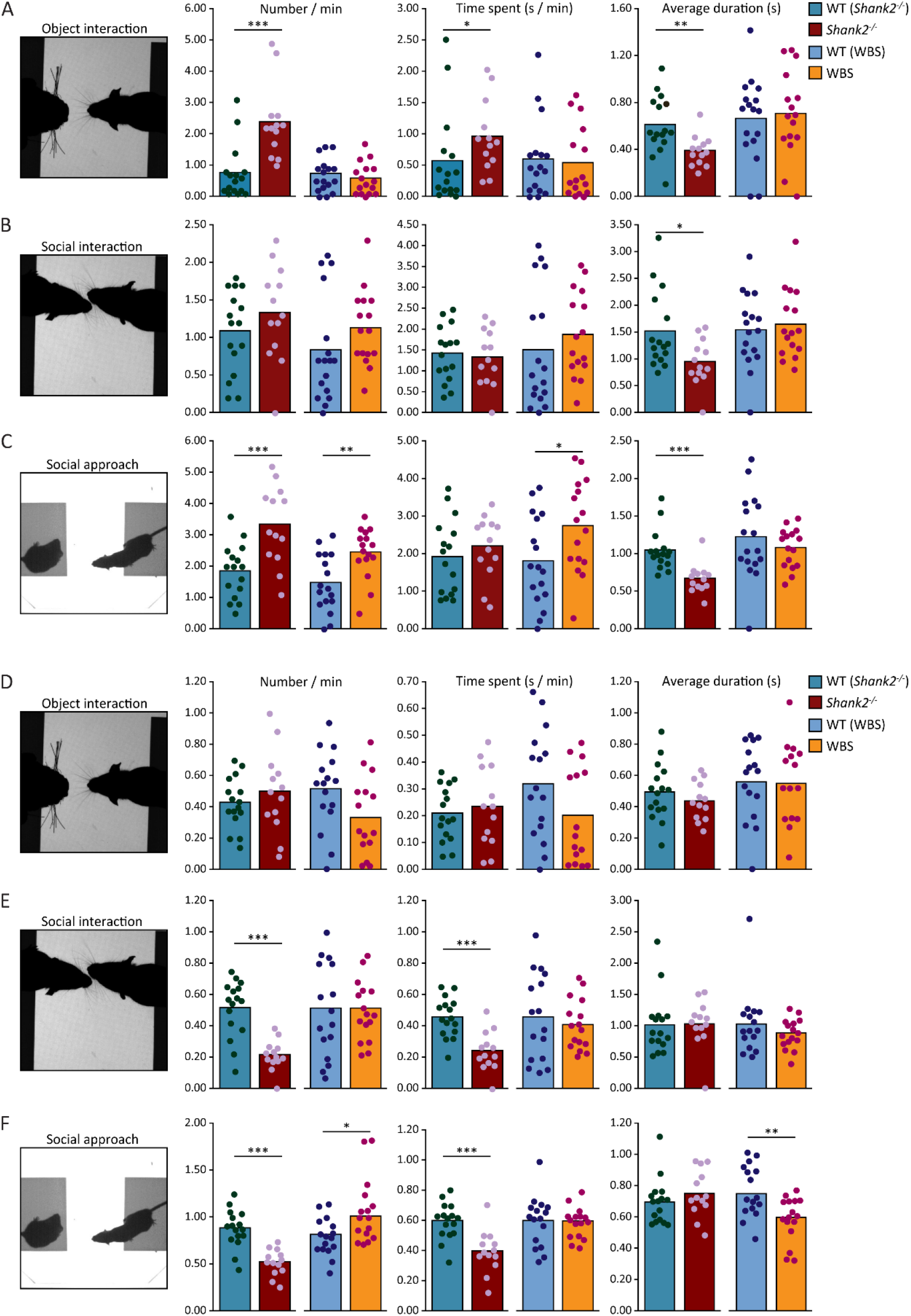
Assessment of phenotypes in social behavior (individual outcome measures underlying the differences in Fig. 5). **A**. Quantification of the number of (left), time spent on (middle) and average duration of (right) object interactions for *Shank2*^-/-^ mice (dark red; N=13), their wild-type litter mates (green blue; N=16), WBS mice (orange; N=16) and their wild-type littermates (blue; N=17). **B**. As in **A**. but for social interactions. **C**. As in **A**. but for social approaches. **D-F**. As in **A-C**. after normalization to gap crossings. For all comparisons: ^*^*p* < 0.05, ^**^*p*<0.01, ^***^*p* < 0.001 (Mann-Whitney U tests).

